# A Novel Central-Peripheral Interface: The Auditory Nerve Glial Transition Zone Exhibits Enhanced Age-Related Immune and Glial Cell Dysfunction

**DOI:** 10.64898/2026.03.27.714751

**Authors:** Shelby A. Payne, Haley R. Anderson, Jiaxin Chai, Peng Chen, Hai Yao, Jeremy L. Barth, Hainan Lang

## Abstract

Age-related hearing loss (ARHL) is a rapidly growing public health concern, affecting two-thirds of adults over 65 years old, with no effective therapeutics available. As the aging population grows at an unprecedented rate, the burden of ARHL will only increase. The causes of ARHL are multifactorial, but an understudied major contributor is glial dysfunction. The auditory nerve (AN) conducts sound from the cochlea to the brainstem and holds a diverse population of immune cells and myelinating glia. As the AN fibers bundle together within the cochlea to project to the brainstem, they are first myelinated by Schwann cells in the peripheral AN, then myelinated by oligodendrocytes in the central AN. The region where myelination shifts from Schwann cells to oligodendrocytes is the glial transition zone (GTZ), located in the cochlear modiolus, creating a unique biological niche. While central-peripheral interfaces are recognized in other cranial nerves, the AN GTZ is understudied. This region integrates the peripheral and central microenvironments within the confined bony cochlea, positioning it as a niche for glial dysfunction in pathological conditions, such as aging. We hypothesize that the GTZ is a site of enhanced glial dysfunction contributing to age-related AN demyelination, an important contributor to ARHL. We evaluated this in an ARHL mouse model combining RNA-sequencing, quantitative immunohistochemistry, and 3D high-resolution imaging. We examined the AN GTZ from human temporal bone donors. RNA-sequencing of the AN revealed age-associated increases in abnormal myelination/glial function and inflammation. There was a significant age-dependent increase in Iba1^+^ macrophages/microglia, with accumulation at the AN GTZ, and an increase in cellular volume and surface area, suggesting greater age-related activation. Macrophages/microglia contained significantly more internalized myelin debris in the AN (peripheral, central, and GTZ) with aging. More importantly, we found structurally intact myelin within macrophages/microglia only at the GTZ, suggesting a unique microenvironment at the GTZ altering phagocytic activity in aging. Together, our data suggest that the GTZ, a previously unrecognized central-peripheral interface, is a critical site of immune-glial interactions and especially vulnerable to age-related demyelination and neuroinflammation. This study highlights the GTZ as a potential target for preserving AN myelination and mitigating ARHL.

## 1. Introduction

Two-thirds of adults over the age of 65 years old are affected by age-related hearing loss (ARHL) (National Center for Health Statistics (U.S.) *et al*., 2021). Although there are numerous contributors to the pathology of ARHL, glial dysfunction remains understudied. The auditory nerve (AN), which is one branch of cranial nerve VIII, conducts sound from the cochlea to the brainstem (Bordoni, Mankowski and Daly, 2025). This nerve also contains a diverse population of immune cells and myelinating glia (Hu, Zhang and Frye, 2018). The peripheral AN is myelinated by Schwann cells, while the central AN is myelinated by oligodendrocytes (Bercury and Macklin, 2015; Long *et al*., 2018). The region where myelination shifts from Schwann cells to oligodendrocytes is called the glial transition zone (GTZ) (i.e. Obersteiner-Redlich zone) and creates a unique biological niche due to the close proximity of both peripheral and central glia (Fontenas, 2023). While such central-peripheral interfaces have been defined in other cranial nerves and the spinal cord, the GTZ is understudied in the auditory system (Fraher, 1992; Knipper *et al*., 1998; Osen, Furness and Hackney, 2011; Bojrab *et al*., 2017; Li *et al*., 2022). The GTZ of the AN is a one-of-a-kind region of glial diversity because it integrates the peripheral and central microenvironments within the bony confines of the cochlea, which is housed in the petrous portion of the temporal bone, the hardest bone in the human body. This makes the GTZ of the AN a niche for concentrated glial dysfunction in pathological conditions, such as aging, which is the focus of this study.

Previous studies in the spinal cord suggest that such central-peripheral transition zones serve as important immune interfaces that regulate myelin turnover during development and after injury (Borjini *et al*., 2019; Wu *et al*., 2022; Fontenas, 2023; Xin *et al*., 2025). Within the brain, both resident microglia and peripheral macrophages play important roles in myelination during development and myelin maintenance, consistent with the dynamic nature of myelination throughout an individual’s lifetime (Chapman and Hill, 2020; Santos and Fields, 2021; Kent and Miron, 2023; Gao *et al*., 2024). However, in the aging nervous system, these processes become dysregulated, leading to myelin breakdown, chronic inflammation, and axonal degeneration (Beirowski *et al*., 2014; Raj *et al*., 2017; Watson *et al*., 2017; Borucki *et al*., 2020; Beirowski, 2022; Seicol, Lin and Xie, 2022; Huang *et al*., 2025). Myelin is critical to AN function because it preserves temporal synchrony required for speech perception (Long *et al*., 2018; Harris *et al*., 2021, 2022; Kister and Kister, 2023). However, the effects of aging on the immune and glial cell populations at the AN GTZ housed within the cochlea have not been defined.

Understanding the dynamics of the AN immune cell compartment and associated changes to myelination within the aging AN is critical for determining the mechanisms of ARHL and other cochlear pathologies. Although prior studies have described macrophages in the cochlea during development, aging, and acute noise injury (Fischer *et al*., 2020; Noble *et al*., 2022; Seicol, Lin and Xie, 2022; Song *et al*., 2022; Lang *et al*., 2023), the effects of aging on the interactions of AN immune cells and myelinating glia remain largely unexplored. Furthermore, the extent to which these age-related changes differ between the central AN, the peripheral AN, and the GTZ has not been addressed in mouse models or in human temporal bones.

The goal of this study is to better understand aging effects on AN myelination and immune-glial interactions, specifically focusing on age-related differences within the peripheral AN, central AN, and GTZ, a unique niche of glial-immune interactions in the context of age-related AN degeneration. Utilizing RNA-sequencing, quantitative immunohistochemistry, and 3D high-resolution imaging in an established ARHL mouse model, we examined aging effects on the AN GTZ, macrophage/microglia populations, myelination, glia-immune interactions, and phagocytic activity within the central AN, the peripheral AN, and the AN GTZ. We also assessed the AN GTZ from human temporal bone donors.

## 2. Materials and Methods

### 2.1 Animals

All studies were performed in accordance with the guidelines of the Institutional Animal Care and Use Committee of the Medical University of South Carolina (MUSC) and the Ralph H. Johnson VA Health Care System (VAHCS) in Charleston, South Carolina. Breeding pairs for CBA/CaJ mice were originally purchased from The Jackson Laboratory (JAX#000654) and bred at the MUSC Animal Research Facility. Mice were bred and housed in a low-noise vivarium with a 12/12-hour light/dark cycle and given standard lab chow and water ad libitum. Young adult (3-4 months) and aged (>2.5 years) CBA/CaJ mice (of both sexes) were used in this study. The number of mice per group is reported in the figure legends. Mice with signs of external ear canal obstruction, middle ear obstruction, or infection were excluded.

### 2.2 Assessment of auditory function

All mice used in this study underwent auditory brainstem recordings (ABRs) as previously described (Brown *et al*., 2017; Panganiban *et al*., 2022; Fabrizio-Stover *et al*., 2025). Mice were anesthetized with an intraperitoneal injection (10 mg/kg xylazine and 100 mg/kg ketamine). ABR recordings were performed in an acoustically isolated booth (IAC Acoustics), and mice were placed on a 37°C heating pad, with artificial tear ointment on both eyes. Subdermal needle electrodes (F-EZ-24, Genuine Grass Reusable Subdermal Needle Electrodes) were placed on the vertex (recording), ipsilateral mastoid (reference), and hind limb (ground). ABRs were digitized at 15 kHz using a low-impedance head stage connected to a pre-amplifier (RA4LI/RA4PA, Tucker Davis Technologies). Electrode impedance was checked at the start of each session to ensure it did not surpass 3 kΩ. The pre-amplifier was connected to an RZ6 input/output device (Tucker Davis Technologies) and responses were recorded with BioSigRZ software (Tucker Davis Technologies). Audiometric thresholds were evaluated with 1.1 ms tone pips with 0.55 ms cosine rise-fall time presented from 90 dB SPL to 5 dB SPL using 5 dB steps. Stimuli were presented in a closed-field through an MF1 speaker (Tucker Davis Technologies) coupled to a 3 mm diameter plastic tube and earpiece inserted into the ear canal. Presentation rate of stimuli was 21 Hz with an average of 512 repetitions. ABR Wave I thresholds at 11.3 kHz were determined via the lowest stimulus level with an identifiable and reproducible Wave I (Supplementary Figure 1).

### 2.3 Mouse cochlear tissue collection and preparation

Mice were euthanized and transcardial perfusion was performed with 4% paraformaldehyde (Electron Microscopy Sciences) solution in 1x phosphate buffered saline, pH 7.4 (1xPBS). Temporal bones were then collected and immediately bath-fixed at 4 °C with 4% paraformaldehyde solution in 1xPBS and perfused via the round and oval windows. Temporal bones were kept in fixative for 24 hours at 4 °C. Fixed cochleae were then decalcified in 10% Ethylenediamine tetraacetic acid (EDTA) for 1-3 days depending on age of the mouse. Each cochlea was then cryoprotected in a 15%/20%/30% sucrose gradient over 2 days. After cryoprotection, cochleae were embedded in Tissue-Tek OCT compound and sectioned at a thickness of 30 μm, then stored at −20°C until needed for staining. The sectioning plane of mouse temporal bones was oriented such that the cutting face was parallel to the AN within the cochlear modiolus to ensure that sections were mid-modiolar to allow optimal viewing of the AN GTZ.

### 2.4 Human cochlear tissue collection and preparation

Procedures for the collection and preparation of human temporal bones have been previously reported (Cunningham *et al*., 2001; Xing *et al*., 2012; Noble *et al*., 2019). All specimens were obtained from the MUSC Hearing Research Program’s temporal bone archive and the MUSC Carroll A. Campbell, Jr. Neuropathology Laboratory Brain Bank. In all cases of human temporal bone collection, written informed consent was obtained from the next-of-kin in accordance with South Carolina laws and regulations. Temporal bone research was approved by the MUSC Institutional Review Board as not human subject research (Pro0030845). No hearing history is available for the human temporal bone (89+ year old female donor) used in this study. After removal of the temporal bone from the skull, scalar perfusion was performed with a 4% solution of paraformaldehyde. Fixation was continued by immersion for at least 48 hours at 4°C. Each temporal bone was then rinsed with 1x PBS and decalcified in 0.35 M EDTA, pH 8.0 for a period of 4-6 weeks as described previously (Cunningham et al., 2000). During this decalcification period, each specimen was trimmed to remove the hard bone encasing the cochlea and vestibule of the inner ear. Once fully decalcified and trimmed, the inner ear portions of the temporal bone were processed for frozen sectioning at a thickness of 12 μm.

### 2.5 Immunohistochemistry on mouse and human cochlear sections

Cochlear sections were air dried with a fan for 30 minutes, then submerged in −20°C acetone for 5 minutes followed by −20°C methanol for 10 minutes. Sections were permeabilized and blocked in donkey serum buffer with Triton X-100 (16% donkey serum, 0.3% Triton X-100, 450 mM NaCl, 20 mM Phosphate Buffer) at room temperature for 3 hours. Primary antibodies were prepared in donkey serum buffer with Triton X-100 and added to the sections, then incubated overnight at 4°C. The next day, sections were washed with wash buffer with Triton X-100 (0.3% Triton X-100, 450 mM NaCl, 20 mM Phosphate Buffer) and the appropriate biotinylated secondary antibodies conjugated with fluorescent avidin (Vector Labs) were added as previously described (Lang et al., 2011). For myelin staining with FluoroMyelin^TM^ (ThermoFisher Scientific), slides were washed and blocked, then stained overnight at 4°C, based on the protocol recommended by the manufacturer. The following day slides were washed using wash buffer with Triton X-100, stained for nuclei (Hoechst), then washed again before being mounted with Vectashield (Vector Laboratories). The primary antibodies, secondary antibodies, and stains used for immunohistochemistry are listed in Table 1.

**Table 1.** Antibodies and stains used for this study.

| **Antibody** | **Source** | **Vendor** | **Catalog Number** | **Dilution** |
| --- | --- | --- | --- | --- |
| Iba1 | Rabbit | Abcam | AB178847 | 1:150 |
| CC-1 | Mouse | Millipore | MABC200 | 1:100 |
| NG2 | Rabbit | Millipore | AB5320 | 1:100 |
| Biotinylated anti-rabbit IgG | Goat | Vector Laboratories | X1104 | 1:100 |
| Biotinylated anti-mouse IgG | Horse | Vector Laboratories | BA2000 | 1:100 |
| Biotinylated anti-rat IgG | Rabbit | Vector Laboratories | BA4000 | 1:100 |
| Streptavidin Texas Red™ | N/A | Vector Laboratories | SA5006 | 1:100 |
| Streptavidin Fluorescein | N/A | Vector Laboratories | SA5001 | 1:100 |
| Anti-mouse IgG Alexa Fluor™ 488 | Rabbit | ThermoFisher Scientific | A11029 | 1:200 |
| Myelin Basic Protein, REAfinity Vio R667 | N/A | Miltenyi Biotec | 130-131-152 | 1:50 |
| CD68, REAfinity Vio G570 | N/A | Miltenyi Biotec | 130-133-803 | 1:50 |
| Toluidine Blue | N/A | Electron Microscopy Sciences | 22050 | 1% |
| FluoroMyelin™ Green | N/A | Invitrogen | F34651 | 1:700 |
| Hoechst 33342 | N/A | ThermoFisher Scientific | H3570 | 1:500 |

### 2.6 Confocal imaging with Airyscan and post-acquisition analysis

Slice and confocal image stacks were collected using a Zeiss LSM 880 NLO with Airyscan using ZEN acquisition software (Zeiss United States). For human cochlear section preparations, images were taken at a resolution of 0.52 µm x 0.52 µm to evaluate the AN and AN GTZ. Imaging was performed at a resolution of 0.16 µm x 0.16 µm to analyze macrophage/microglia morphology and myelination in human temporal bone sections. For mouse cochlear section preparations, images were taken at a resolution of 0.83 µm x 0.83 µm for evaluating the AN and AN GTZ. For analysis of macrophage/microglia morphology and myelination (including assessment of internalized myelin), image stacks were taken at a resolution of 0.13 µm x 0.13 µm with a z-step of 0.60 µm. Images were processed using ZEN 3.9 Blue Edition (Carl Zeiss Microscopy GmbH).

The Imaris volume rendering function (Imaris 10.2, Oxford Instruments) was used for 3D reconstructions and analysis of macrophage/microglia morphology, myelination, and identification and quantification of internalized myelin within AN macrophages/microglia.

Immunofluorescence image stacks from 30 µm sections in CBA/CaJ mice of both age groups were imported into Imaris 10.2. For the 3D reconstruction of Iba1^+^ cells, surfaces were created using the Machine Learning Segmentation Tool (Imaris 10.2, Oxford Instruments). Training was performed on each image using 3-5 repetitions as needed, based on complexity of macrophage/microglia morphology. Any touching objects were split. Surfaces were hand-filtered to remove any objects that were not part of the Iba1^+^ macrophage/microglia mask. All surface parameters were exported into a separate .csv file for each image stack.

Myelination was quantified in the same manner, using the Machine Learning Segmentation Tool (Imaris 10.2, Oxford Instruments), which generated surfaces of each myelinated fiber in the section and output the surface area and volume of the myelinated fibers contained within each region of interest (ROI). For the 3D reconstruction of myelinated fibers in the AN to evaluate surface area and volume, the same procedure was used as described above for generating surfaces of Iba1^+^ macrophages/microglia. After the surfaces of the myelinated fibers were created for each image, the statistics were exported, including the total volume and surface area of each surface. All surface parameters were exported into a separate .csv file for each image stack.

The quantity of internalized myelin was determined based on the volume of the myelin surface contained within the Iba1^+^ surface to ensure only fully enclosed myelin components were included in the analysis. For quantification of total internalized myelin within Iba1^+^ macrophages/microglia, Object-Object Statistics were performed in Imaris (Imaris 10.2, Oxford Instruments). For each individual image, the Iba1^+^ macrophage/microglia surface was used to create a mask against the myelinated fiber surface generated for each stack. Then, a surface was created that contained only myelin-stained objects within the Iba1^+^ macrophage/microglia surface. The statistics for the internalized myelin surface were then exported to determine total volume of internalized myelin within the Iba1^+^ macrophages/microglia for each image stack.

### 2.7 Epoxy resin embedding preparation of the auditory nerve

CBA/CaJ mice were processed and imaged via electron microscopy as previously described (Lang *et al*., 2011). Briefly, mice (CBA/CaJ 1 year old) were perfused via cardiac perfusion with 10 mL normal saline with 0.1% sodium nitrate, then 15 mL of 4% paraformaldehyde and 2% glutaraldehyde in 0.1 M phosphate buffer, pH 7.4. Cochleae were decalcified in 50 mL of 120 mM solution of ethylenediaminetetraacetic acid (EDTA), pH 7.2, with gentle stirring at room temperature for 2-3 days with daily changes of EDTA solution. Tissues were post-fixed in 1% osmium tetroxide for 1 hour, dehydrated, and embedded in Epon LX 112 resin. Semi-thin sections were cut at approximately 1 µm thick and stained with Toluidine blue.

### 2.8 Light sheet microscopy and three-dimensional high-resolution imaging of the auditory nerve glial transition zone

Young and aged mouse temporal bones were tissue-cleared and imaged with large-field light sheet microscopy to visualize the AN GTZ. Tissue clearing was performed using the MACS Clearing Kit (Miltenyi Biotec) and their standard protocol with the addition of a decalcification step (10% EDTA for 1 day) before permeabilization. Mouse temporal bones were stained with CD68 (Miltenyi Biotec) and Myelin Basic Protein (Miltenyi Biotec). Detailed descriptions of antibodies used are listed in Table 1. Cleared whole mouse temporal bones were mounted and imaged with an Ultramicroscopy II Scope (Miltenyi Biotec) with a Super Plan configuration, equipped with a sC-MOS camera (4.2 Megapixel, Andor Technology), and objective lenses with dipping caps specifically designed and optimized for large-field light sheet imaging as previously described (Chen *et al*., 2025). All 3D image stacks were acquired using a 561 nm laser with a 595/40 nm emission filter to capture CD68 fluorescence, and a 639 nm laser with a 680/30 nm emission filter to capture Myelin Basic Protein fluorescence. Each mouse temporal bone was mounted on the sample holder and oriented such that volume was acquired with the z-steps moving from apex to base through the AN to ensure the capture of the full AN GTZ. The entire AN GTZ was imaged using a 12x objective at 0.6x magnification with a resolution of 0.903 x 0.903 (X,Y) with a 1 µm z-step interval. The number of CD68^+^ cells was quantified every 25 µm within the AN for each temporal bone in young and aged CBA/CaJ mice.

### 2.9 RNA-sequencing analysis of the auditory nerves in young and aged mice

To examine expression of genes relating to myelination, glial cell function, and abnormal inflammation in young and aged mouse AN, analysis was performed on bulk RNA-sequencing data obtained from a prior study involving young adult (3 months) and aged (>2.5 years) ANs from CBA/CaJ mice (accession GSE141865) (Panganiban *et al*., 2022). AN tissues in the previous study were isolated by microdissection of the modiolus from the other structures of the cochlear bulla (such as the cochlear lateral wall and sensory epithelium), with the tissue collected from the two cochleae of each mouse pooled to make each biological replicate; three biological replicates of each age group were used. Raw sequencing data (fastq files) were analyzed using Partek Flow software (Illumina Inc). Reads were aligned to the mouse genome assembly mm39 by STAR (version 2.7.8a) (Dobin *et al*., 2013) and quantified to annotation model (Partek E/M) built from mm39 Ensembl Transcripts release 104 using default parameter settings (Strict paired-end compatibility=true; Require junction reads to match introns=true; Minimum read overlap with feature=100%). Normalization and comparative analysis was done with DESeq2 (Love, Huber and Anders, 2014). Normalized count data is archived in NCBI Gene Expression Omibus (accession GSE320402). Differential gene expression was defined as adjusted *p*-value (FDR step up) < 0.05 and absolute fold change > 2, yielding 1240 genes (Supplementary Table 1). Biological process enrichment analysis was conducted with ToppGene (Chen *et al*., 2009).

### 2.10 Statistical analysis

Sample sizes for experiments and morphological observations are listed in the appropriate results section and figure legends. Images presented here are representative of each experimental group for fluorescence staining. All statistical values are presented as a mean ± standard deviation unless stated otherwise. All data sets were tested for normality using a Kolmogorov-Smirnov test with reference to a normal distribution. Mann-Whitney U-test was used for pairwise comparisons between groups. Exact *p*-values are reported for all comparisons. For differential expression analyses of RNA-sequencing data, a *p*-adjusted value (FDR step up) of < 0.05 and absolute fold change > 2 was considered significant. Graphs were plotted and statistics were performed in Igor Pro 9 (Wavemetrics) and GraphPad Prism 8 (GraphPad Software).

## 3. Results

Previous studies on aging effects on neural degeneration of the auditory system have focused on the peripheral AN fibers in the region where they project to the sensory epithelia, specifically on the spiral ganglion and ribbon synapses within the inner hair cells (Sergeyenko *et al*., 2013; Wan and Corfas, 2017; Long *et al*., 2018; Heeringa *et al*., 2020, 2020; Budak *et al*., 2021; Panganiban *et al*., 2022). Other studies have focused on the central auditory system, including the cochlear nucleus and auditory cortex (Cramer and Rubel, 2016; Milinkeviciute *et al*., 2019; Eggink *et al*., 2022; Seicol, Lin and Xie, 2022). Here, this study investigated aging effects on AN myelination and inflammation, focusing on the GTZ which houses both peripheral and central immune cells and myelinating glia, in an established ARHL mouse model and human temporal bone sections from an older donor.

### 3.1 The glial transition zone of the mouse auditory nerve

To better evaluate the interactions between macrophages/microglia and the myelinating glia of the AN GTZ, we used FluoroMyelin^TM^ stain on 30 µm frozen sections of young and aged CBA/CaJ mice (Figure 1A, D; see Table 1 for detailed description). Auditory function was assessed for mice in this study, and aged mice had elevated hearing thresholds compared to young mice (Supplementary Figure 1). Sections were also stained with the primary antibody Iba1 as well as a nuclear marker (Hoechst) to aid in the identification of the cell bodies of Iba1^+^ macrophages/microglia. Three labeling approaches utilized in this study reveal the GTZ, identify a gap in myelin-related structures between central and peripheral portions of the mouse AN (indicated by the dashed white lines), and distinguish the peripheral AN from central AN (Figure 1; Supplementary Figure 2). These labeling approaches include 1) frozen sections stained with FluoroMyelin^TM^ stain (Figure 1A,D), 2) Epon LX 112 resin embedded sections stained with Toluidine blue (Figure 1G, G’), and 3) frozen sections stained with NG2 antibody for Schwann cells (peripheral AN) and CC1 and NG2 antibodies for oligodendrocytes (central AN) (Figure 1C, F; Supplementary Figure 2). FluoroMyelin^TM^ staining revealed region-specific structural differences between the central and peripheral AN. Central AN myelin has a different depth of staining compared to peripheral AN myelin (Figure 1A, D) and appears to have a lower nuclei density than that of the peripheral AN (Figure 1C, F). This observation of increased central myelin staining is consistent with known compositional differences between central and peripheral myelin. Previous studies report that myelin basic protein makes up approximately 30% of the protein mass of central myelin, while it only accounts for 5-18% of the protein mass of peripheral myelin (Garbay *et al*., 2000; Kister and Kister, 2023). Like other cranial nerve transition zones, the higher nuclear density seen in the peripheral AN compared to the central AN reflects the differences in myelinating cell types across the GTZ. This is because the central AN is myelinated by oligodendrocytes, which surround numerous AN fibers with one cell body, while the peripheral AN is myelinated by Schwann cells, which surround only one individual FR. Iba1 staining showed macrophages and microglia dispersed throughout the AN in young and aged sections (Figure 1B, E). In these low-magnification mid-modiolar sections, observable differences in the Iba1^+^ cell populations throughout the AN and at the GTZ are apparent, with the aged sections having a greater number of Iba1^+^ cells throughout the AN and Iba1^+^ cells appearing to accumulate at the GTZ (Figure 1B, E).

**Figure 1.**
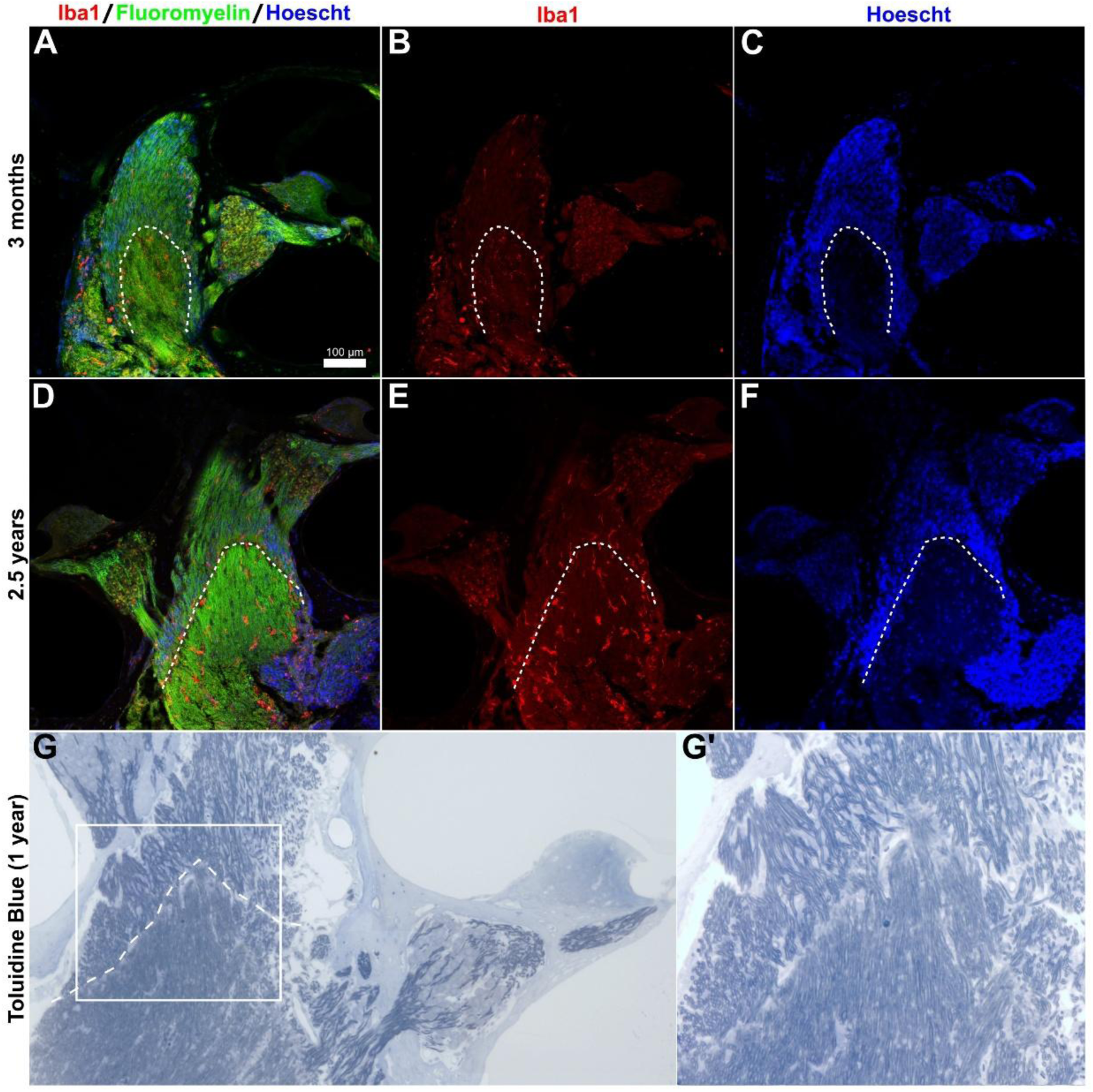
Mouse mid-modiolar sections demonstrating age-related changes to the AN GTZ. **(A-F)** Thirty µm mid-modiolar sections of the auditory nerve (AN) glial transition zone (GTZ) in a young (3 months) (A-C) and an aged (2.5 years) (D-F) CBA/CaJ mouse. Sections are stained with FluoroMyelin^TM^ (green), macrophage/microglia cell marker Iba1 (red), and Hoechst nuclear stain (blue). The dashed white line indicates where the AN GTZ is located in a mid-modiolar section. Scale bar = 100 µm. **(G)** Representative image from Epon LX 112 resin-embedded sections (stained with Toluidine blue) of the AN GTZ in a 1 year-old mouse. **(G’)** Enlarged images of the white outlined region of the AN GTZ are shown to illustrate the visible gap between central and peripheral myelination in the AN.

To examine the entire AN and the GTZ, a temporal bone from a young mouse and an aged mouse were cleared and imaged via large-field light sheet microscopy (Power and Huisken, 2017; Chen *et al*., 2025). Temporal bones were immunostained with a fluorescently tagged Myelin Basic Protein antibody, a reliable marker of AN myelin sheaths (Xing *et al*., 2012). Examining young and aged mouse temporal bones shows that the GTZ spans the entirety of the AN and resides within the cochlear modiolus in a cone-shaped morphology (Figure 2E’, P’). A visible transition from peripheral AN to central AN is again evident based on the differences in Myelin Basic Protein staining intensity throughout the AN in both the young and aged samples. To identify phagocytic and active immune cell subpopulations, temporal bones were also stained with a fluorescently tagged CD68 antibody (Hendrickx *et al*., 2017; Choudhary and Malek, 2023; Swanson *et al*., 2023). CD68^+^ cells were observed more frequently within the AN in the aged temporal bones compared to the young temporal bones, mostly within the central AN as well as accumulating near the AN GTZ (Figure 2T’, V’).

**Figure 2.**
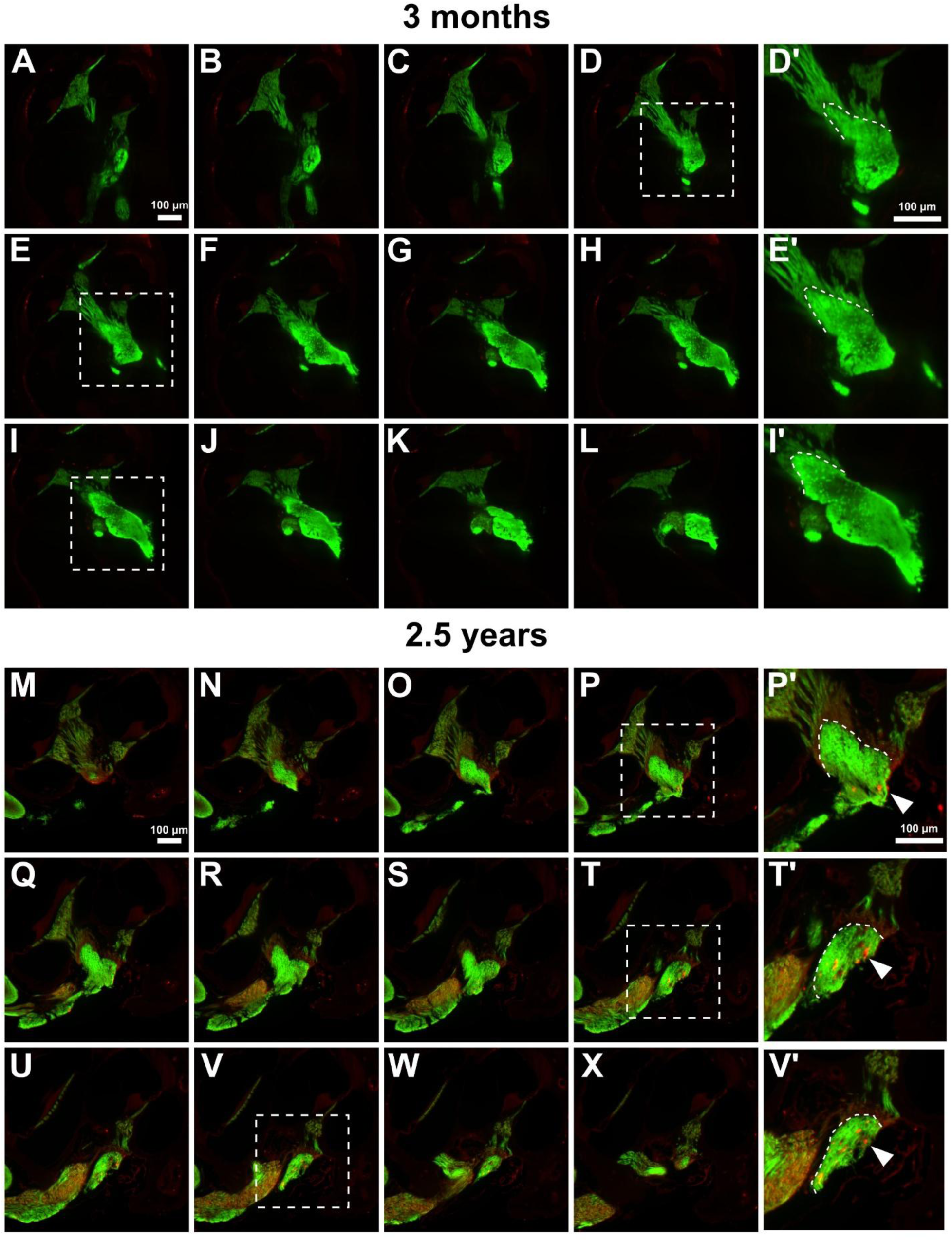
Stacked serial sections of the mouse AN GTZ identifying more CD68^+^ cells in aged mice. **(A –X)** Serial sections taken 25 μm apart from tissue cleared mouse temporal bones showing the entire AN within the cochlea and the AN GTZ in a young (3 months) (A-L) and an aged (2.5 years) (M-X) CBA/CaJ mouse. **(D’, E’, I’)** Enlarged images of the white outlined regions of the AN GTZ in a young mouse are shown. **(P’, T’, V’)** Enlarged images of the white outlined regions of the AN GTZ in an aged mouse are shown with a white arrow indicating CD68^+^ cells within the AN, while no CD68^+^ cells were seen in the AN of the young mouse (D’, E’, I’). Dashed white lines in D’, E’, I’, and P’ show the cone-shaped morphology of the AN GTZ. Dashed white lines in T’ and V’ show the GTZ in the basal turn/hook of the cochlea. Each mouse temporal bone was stained with Myelin Basic Protein (green) and CD68 (red). Scale bars = 200 µm for all images.

### 3.2 Aging auditory nerves have more immune cells, with focal accumulation at the glial transition zone

Previous studies have reported age-related changes to cochlear macrophages/microglia within the spiral ganglia and cochlea (Noble *et al*., 2019; Fischer *et al*., 2020). To assess the effects of aging on AN macrophages/microglia at the AN GTZ, we performed quantitative immunohistochemistry to evaluate Iba1^+^ cells by region of the AN. Immunostained sections of the peripheral AN, GTZ, and central AN are shown in young (Figure 3A-F) and aged (Figure 3G-L) mice.

**Figure 3.**
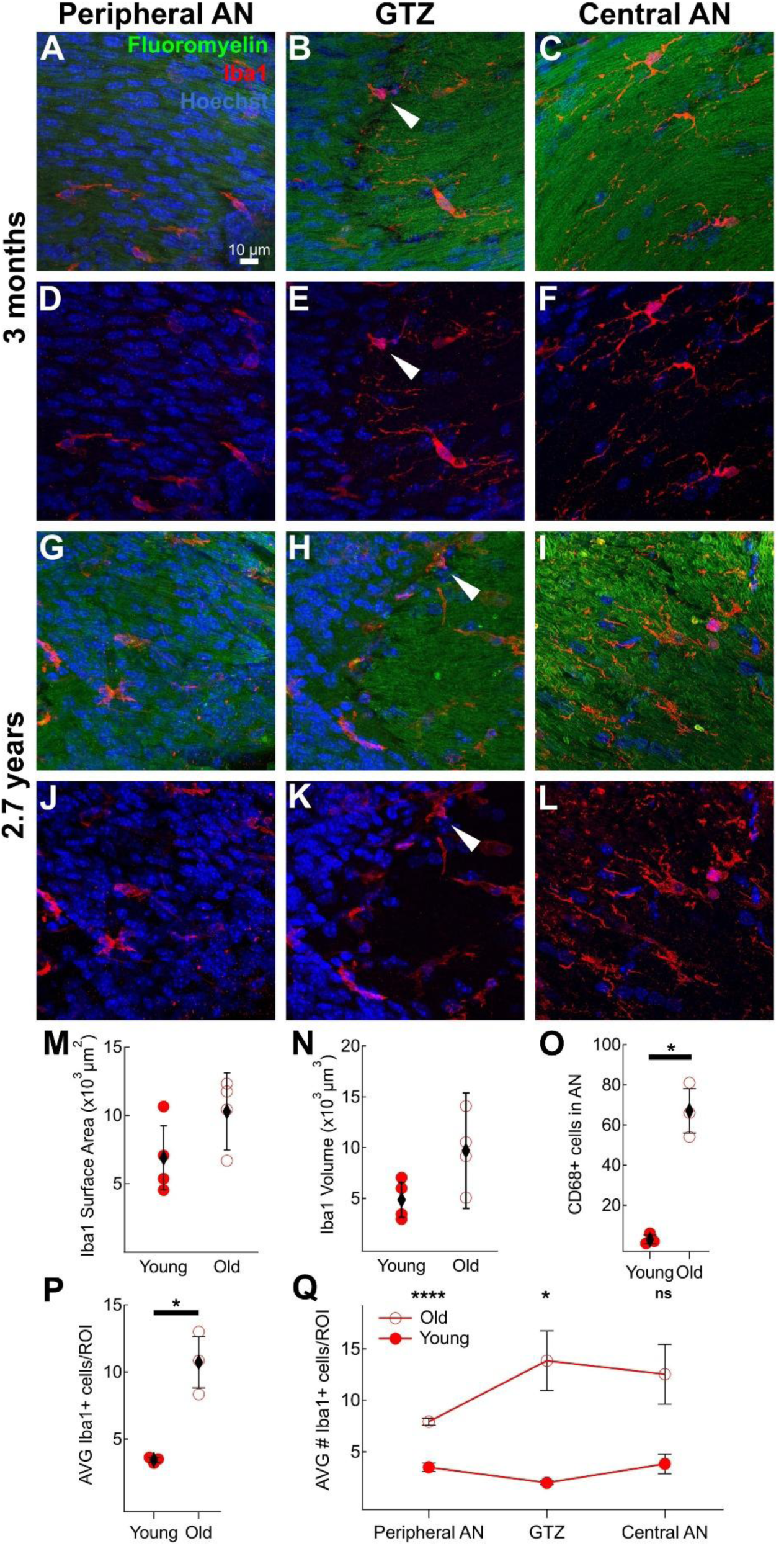
Increased density and morphological alterations of immune cells at the AN GTZ in aged mice. **(A-L)** Immunostained sections of the AN GTZ in young (A-F) and aged (G-L) mice showing Iba1^+^ cells in the peripheral AN (A, D, G, J), GTZ (B, E, H, K), and the central AN (C, F, I, L) with aging. White arrows indicated Iba1^+^ cells that cross both the peripheral and central AN. **(M)** Average surface area of Iba1^+^ cells per region of interest (ROI) in young (filled red circles) and aged (open red circles) mice. N = 4 mice per group. 4 ROIs per mouse (1 ROI for peripheral AN, 1 ROI for central AN, 2 ROIs for AN GTZ). **(N)** Average volume of Iba1^+^ cells per ROI per age group. N = 4 per group. 4 ROIs per animal (1 ROI for peripheral AN, 1 ROI for central AN, 2 ROIs for AN GTZ). **(O)** Quantity of CD68^+^ cells within the entire AN in every section, 25 µm apart in young mice (n=3) and aged mice (n=3). **(P)** Average number of Iba1^+^ cells per ROI in young and aged mice. N = 3 per group, 9 ROIs (3 ROIs for peripheral AN, 3 ROIs for central AN, 3 ROIs for AN GTZ) per mouse. The difference between young and old was significant (*p = 0.03*). **(Q)** Average number of Iba1^+^ cells in the peripheral AN, GTZ, and central AN in young and aged mice. N = 3 mice per group, 3 ROIs per region per age group. All distributions were evaluated for normality using a Kolmogorov-Smirnoff test. Significance was determined using a Mann-Whitney U-test, α = 0.05. Diamonds show the mean, and error bars represent the standard deviation. \**p<0.05*. \*\*\*\**p<0.0001*.

To evaluate the activation state of Iba1^+^ cells within the AN, we assessed their morphology, including surface area and volume (Kopper *et al*., 2021; You *et al*., 2023). Generally, Iba1^+^ cells in aged mice seem to have greater surface areas and volumes compared to young, however, there is no significant difference between groups (Surface Area: AVG_Young_ = 6.90 ± 2.35 x 10^3^ µm^2^, AVG_Old_ = 10.29 ± 2.82 x 10^3^ µm^2^, *p* = 0.111; Volume: AVG_Young_ = 4.86 ± 1.71 x 10^3^ µm^3^, AVG_Old_ = 9.71 ± 5.67 x 10^3^ µm^3^, *p* =0.064) (Figure 3M, N).

To assess phagocytic activity of the Iba1^+^ cells that populate the AN, we stained for CD68, an established marker of phagocytic activation, in the AN with aging (Hendrickx *et al*., 2017; Choudhary and Malek, 2023; Swanson *et al*., 2023). Iba1 acts as a marker of cells of myeloid lineage, such as macrophages and microglia, while CD68 is marker of phagocytic activity by macrophages and microglia(Schwabenland *et al*., 2021). Using serial sections extracted from the cleared mouse temporal bones, we quantified the number of CD68^+^ cells within the entire AN in young mice (3 months) and aged mice (2.5 years) (Figure 2). The aged mice had a significantly greater quantity of CD68^+^ cells compared to the young mice (AVG_Young_ = 3.00 ± 2.16, AVG_old_ = 67.00 ± 11.05, *p* = 0.025) (Figure 3O).

Aged mice contained significantly more Iba1^+^ cells compared to young mice across all regions of the AN (AVG_Young_ = 3.44 ± 0.18, AVG_Old_ = 10.72 ± 1.91, *p* = 0.032) (Figure 3P). Of all three regions of the AN, the GTZ had the greatest density of Iba1^+^ cells in aged mice compared to young mice (Peripheral AN: AVG_Young_ = 3.50 ± 0.41, AVG_Old_ = 7.93 ± 0.32, *p* = 3.67E-04; GTZ: AVG_Young_ = 3.89 ± 0.16, AVG_Old_ = 12.52 ± 2.90, *p* = 0.050; Central AN: AVG_Young_ = 1.67 ± 0.94, AVG_Old_ = 12.57 ± 2.90, *p* = 0.019) (Figure 3Q). Iba1^+^ cells are still present in young mice but sparsely populate the AN in both the peripheral and central regions. Interestingly, Iba1^+^ cells in the AN GTZ were also found with projections in the central and peripheral regions of the AN (Figure 3B, E, H, K).

### 3.3 Aging is associated with disruption of auditory nerve myelin

Myelin staining revealed a difference in morphology between the peripheral AN and central AN in young and aged mice (Figure 4). The peripheral AN had a homogenous myelin stain, while the central AN had a more heterogeneous appearance in both young and aged mice. We observed an increase in disrupted myelin sheaths throughout the peripheral and central AN regions with age (Figure 4A, B, D, E), which confirms previous electron microscopy studies in the peripheral AN (Xing *et al*., 2012; Panganiban *et al*., 2022). To quantify age-related changes in myelination in the AN, we measured the volume and surface area of myelinated fibers in young (3 - 4 months) and aged (≥ 2.5 years) CBA/CaJ mice using Imaris. Aged mice had no significant difference in surface area of myelinated fibers compared to young mice (Figure 4C). Aged mice also had no significant difference in volume of myelinated fibers compared to young mice (Figure 4F). Although not significant, young mice have a larger average surface area and volume of myelinated fibers in the AN compared to aged mice (Surface Area: AVG_Young_ = 183.75 ± 61.00 x 10^2^ µm^2^, AVG_Old_ = 133.74 ± 38.30 x 10^2^ µm^2^, *p* = 0.342; Volume: AVG_Young_ = 290.74 ± 52.63 x 10^3^ µm^2^, AVG_Old_ = 226.50 ± 60.45 x 10^3^ µm^2^, *p* = 0.200).

**Figure 4.**
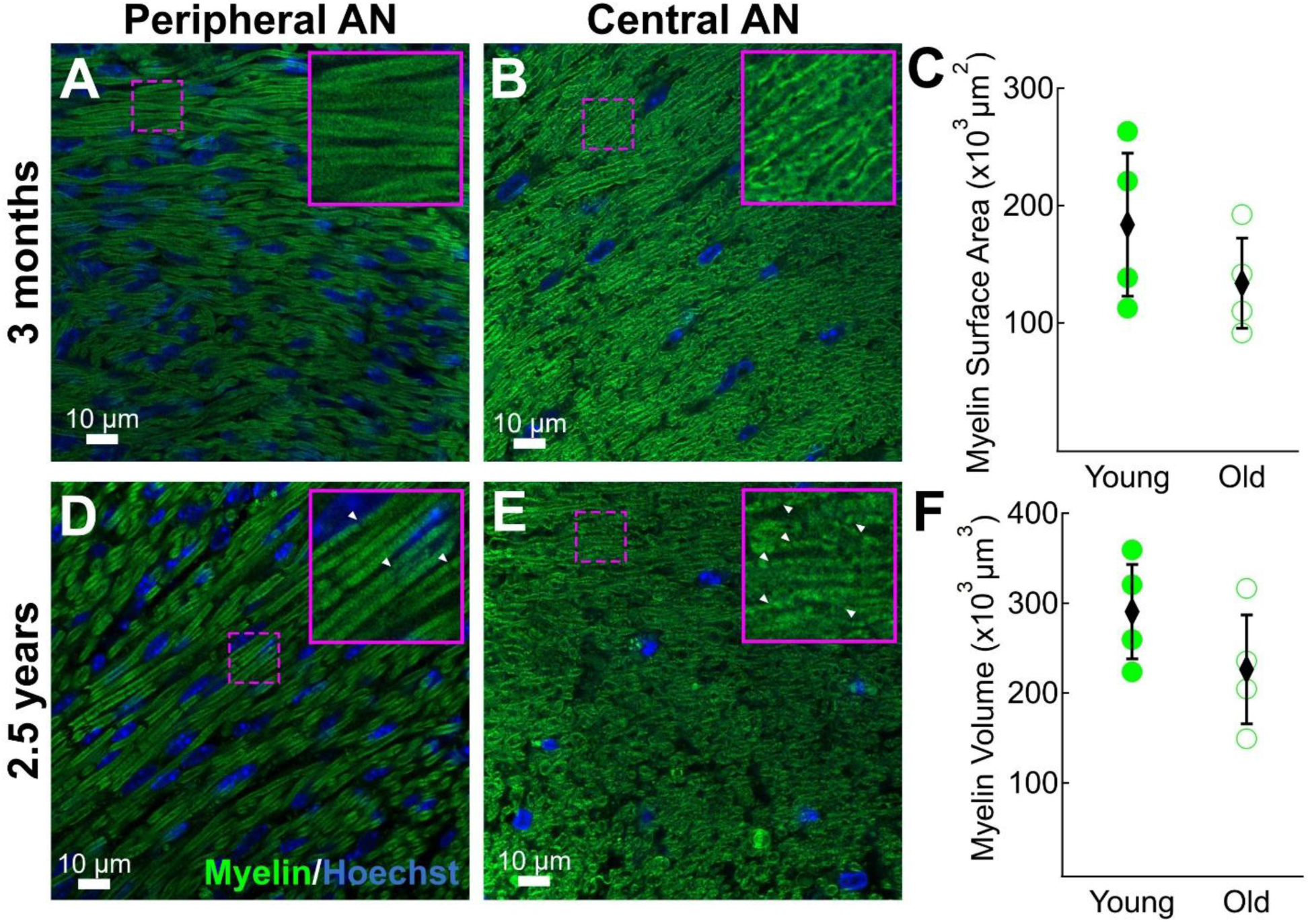
No significant difference in myelin surface area and volume, but more disrupted myelin sheaths in the peripheral AN and central AN in aged mice. **(A-B)** FluoroMyelin^TM^-stained sections in the peripheral AN (A) and central AN (B) in a young (3 months) and aged (2.5 years) CBA/CaJ mouse. Boxes on the top-right are enlarged images from the dashed boxes in A and B. **(C)** Quantification of myelinated AN fiber surface area in young (filled green circles) and aged (open green circles). N= 4 mice per age group. 4 ROIs per mouse (2 ROIs for peripheral AN, 2 ROIs for central AN). The difference between the young and the aged was not significant. **(D-E)** FluoroMyelin^TM^-stained sections in the peripheral AN (D) and central AN (E) in an aged CBA/CaJ mice. Magenta dashed boxes indicate the region of the pop-out image in the magenta outlined box. Boxes on the top-right are enlarged images from the dashed boxes in D and E. White arrows indicate the disrupted myelin sheaths in the peripheral AN (D) and central AN (E). **(F)** Quantification of myelinated AN fiber volume by age group. N = 4 mice per age group. 4 ROIs per animal. The difference between the young and the aged was not significant. Diamonds represent the mean and error bars represent the standard deviation for each group. All distributions were tested for normality using a Kolmogorov-Smirnoff test. Significance was determined using a Mann-Whitney U-test, α = 0.05. Scale bar = 10 µm for all images.

### 3.4 Aging effects on the auditory nerve transcriptome – abnormal immune cell and myelinating glial function

To further validate our observations of age-related myelin degeneration and increased immune cell activity within the AN and its enhanced effects at the AN GTZ, bulk RNA-sequencing data obtained from a prior study of AN from young adult mice (3 months) and aged mice (2.5 years) was analyzed anew, using updated reference databases. Samples in the study contained central AN and peripheral AN within the cochlear modiolus and internal auditory meatus. Analysis of the RNA-sequencing data revealed significant age-dependent transcriptional changes in 1240 genes (FDR < 0.05; absolute fold change > 2) (Figure 5A). Functional enrichment analysis of these genes detected significant effects on pathways relating to myelination and immune activation, including abnormal myelination (q = 2.30 x 10^-2^) and abnormal inflammatory response (q = 1.76 x 10^-9^) (Figure 5B, C). Significantly upregulated genes mapping to these pathways included *Clec7a*, *Trem2*, and *Itgb2* (abnormal inflammatory response). A complete list of differentially expressed genes, and their association(s) with these pathways, is given in Supplementary Table 1. Detection of an effect on abnormal myelination aligns with our observed immunohistochemistry results in old AN. Together, these findings demonstrate that aging is associated with dysregulation of myelin-associated processes and inflammatory signaling.

**Figure 5.**
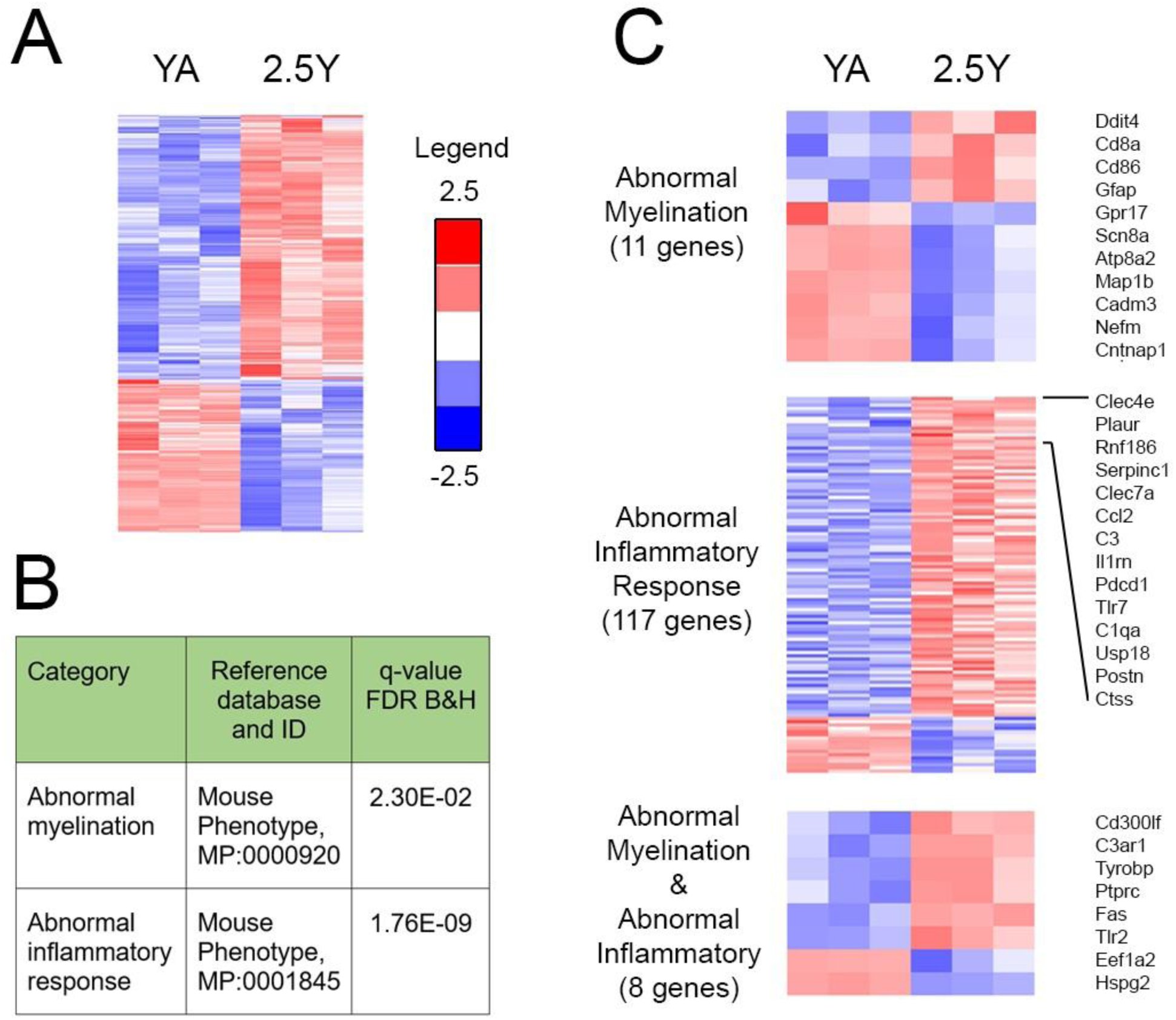
Gene expression profiles of the AN revealed age-dependent effects on inflammation and myelination. **(A)** Analysis of AN RNA-sequencing data from young (3 months) and aged (2.5 years) CBA/CaJ mice detects 1240 differentially expressed genes (Supplementary Table 1). **(B)** Enrichment analysis of differentially expressed genes detects significant effects on abnormal myelination and abnormal inflammatory response. **(C)** Expression patterns of differentially expressed genes linked to abnormal myelination, abnormal inflammatory response or both categories. For abnormal inflammatory response, the 15 genes showing highest upregulation are listed on the right.

### 3.5 Accumulation of myelin debris found in aging auditory nerve macrophages/microglia

Given the proven function of macrophages/microglia in phagocytic processes associated with myelin maintenance, we evaluated this process in aging Iba1^+^ cells (Benmamar-Badel, Owens and Wlodarczyk, 2020; Borucki *et al*., 2020; Hughes and Appel, 2020; Goddery *et al*., 2021; Santos and Fields, 2021; Kent and Miron, 2023, 2023; Swanson *et al*., 2023; Beiter, Sheehan and Schafer, 2024; Berglund *et al*., 2024). Upon close examination of the Iba1^+^ cells within the AN, many contained internalized myelin debris, especially in the aged ANs (Figure 6). The increased presence of Iba1^+^ cells burdened with internalized myelin suggests possible immune dysfunction and dysregulation of the myelination process (Moreno-García *et al*., 2018; Burns *et al*., 2020; Swanson *et al*., 2023; Beiter, Sheehan and Schafer, 2024). Young mice had very little to no internalized myelin debris within Iba1^+^ cells (Figure 6A-C), while aged mice had large amounts of internalized myelin debris within their cell bodies (Figure 6D-F). This is evident in the immunofluorescence images, as well as in the Imaris reconstructions (Figure 6D’, B’). These age-related differences were consistent across the peripheral, central, and GTZ regions of the AN. Quantifying the volume of internalized myelin contained within Iba1^+^ cells, aged mice had significantly more internalized myelin compared to young mice (Internalized Myelin Volume: AVG_Young_ = 9.80 ± 10.69 µm^3^, AVG_Old_ = 686.71 ± 414.09 µm^3^, *p* = 0.004) (Figure 6H). To further explore the effects of aging on phagocytic functions, our bulk RNA-sequencing results were re-examined with the finding that phagocytic categories were significantly enriched in the differentially expressed gene set, including “phagocytosis, engulfment” (GO:0006911; q = 3.67 x 10^-4^). Review of expression patterns for phagocytic engulfment genes illustrated that they were predominantly upregulated in aged AN (Figure 6G).

**Figure 6.**
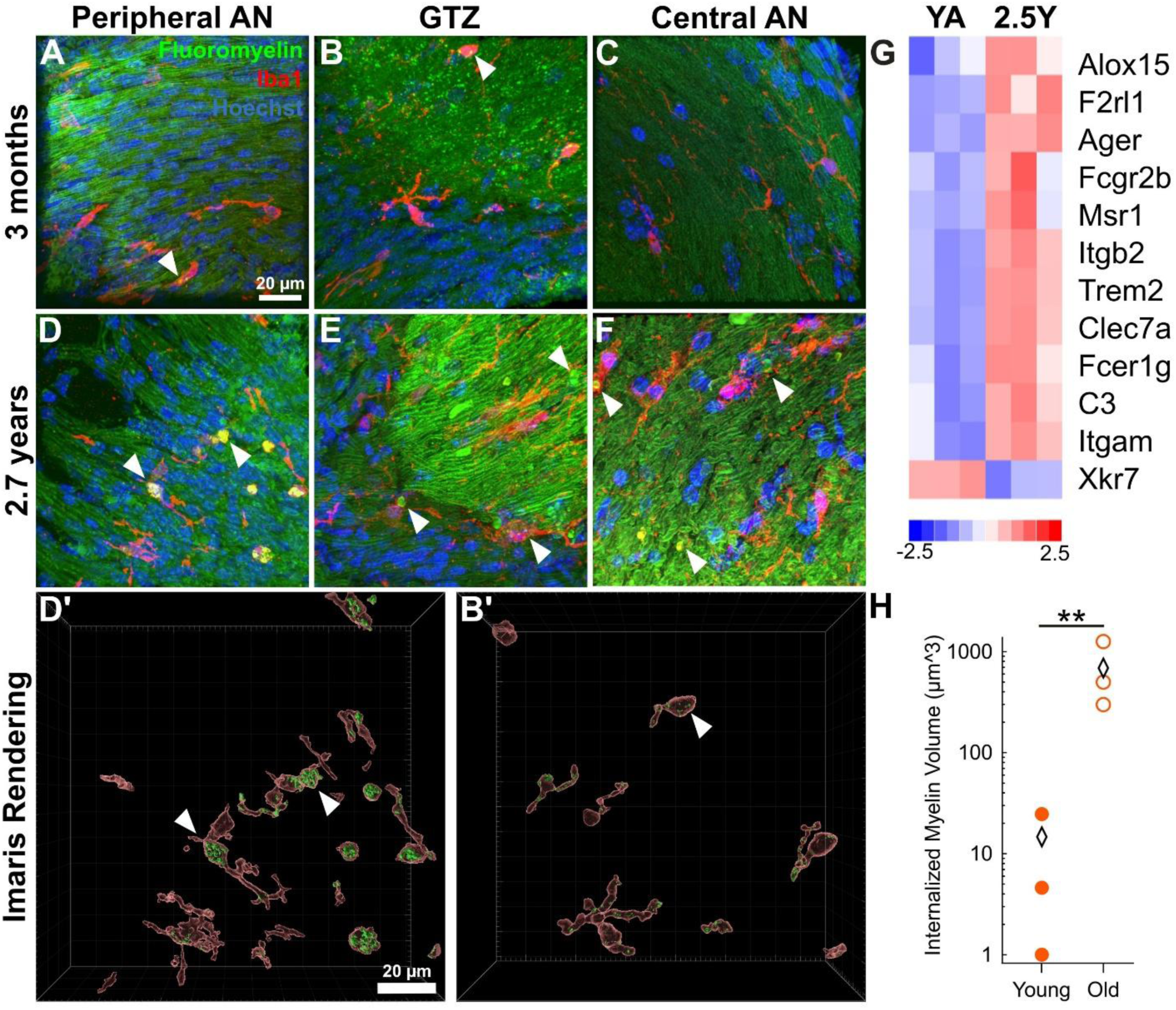
Increased internalized myelin debris and upregulated expression of phagocytosis-related genes appeared in the aged AN. **(A-F)** Sections of the peripheral AN (A, D), GTZ (B, E), central AN (C, F) and Imaris 3D reconstructions (D’, B’) showing myelin debris inside Iba1^+^ cells. **(G)** Differentially expressed genes linked to phagocytic engulfment are predominantly upregulated in aged (2.5 years) versus young (3 months). **(H)** Volume of internalized myelin in young mice and aged mice. 4 ROIs per mouse, N = 3 mice per age group. All distributions were assessed for normality using a Kolmogorov-Smirnoff test. Comparisons between young and old for significance were determined using a Mann-Whitney U-test, α = 0.05. \*\**p < 0.01*.

### 3.6 Iba1^+^ cells at the auditory nerve glial transition zone contain healthy-like myelin

Examining morphological differences in the internalized myelin within Iba1^+^ macrophages/microglia in the peripheral AN, central AN, and the AN GTZ reveals a unique morphology of internalized myelin found only in the aging AN GTZ. Young mice had very few Iba1^+^ cells with any internalized myelin across regions of the AN, including the AN GTZ (Figure 7A-G). By contrast, aged mice had numerous Iba1^+^ cells with internalized myelin debris throughout the AN, but the AN GTZ macrophages/microglia contained myelin that appeared healthy and structurally intact (Figure 7H-N). Upon close inspection of the higher magnification images, there are pockets within the Iba1^+^ cell bodies containing intensely FluoroMyelin^TM^-stained myelin structures, which have a similar morphology to the healthy myelinated fibers in these sections (Figure 7I-N). Additionally, the morphology of this healthy-like myelin has the appearance of an open cylinder, which is different from the disorganized appearance of myelin debris shown in Figure 6. These striking observations suggest that the phagocytic processes performed by immune cells at the GTZ involves both myelin debris and healthy-like, structurally intact myelin sheaths of aged AN fibers.

**Figure 7.**
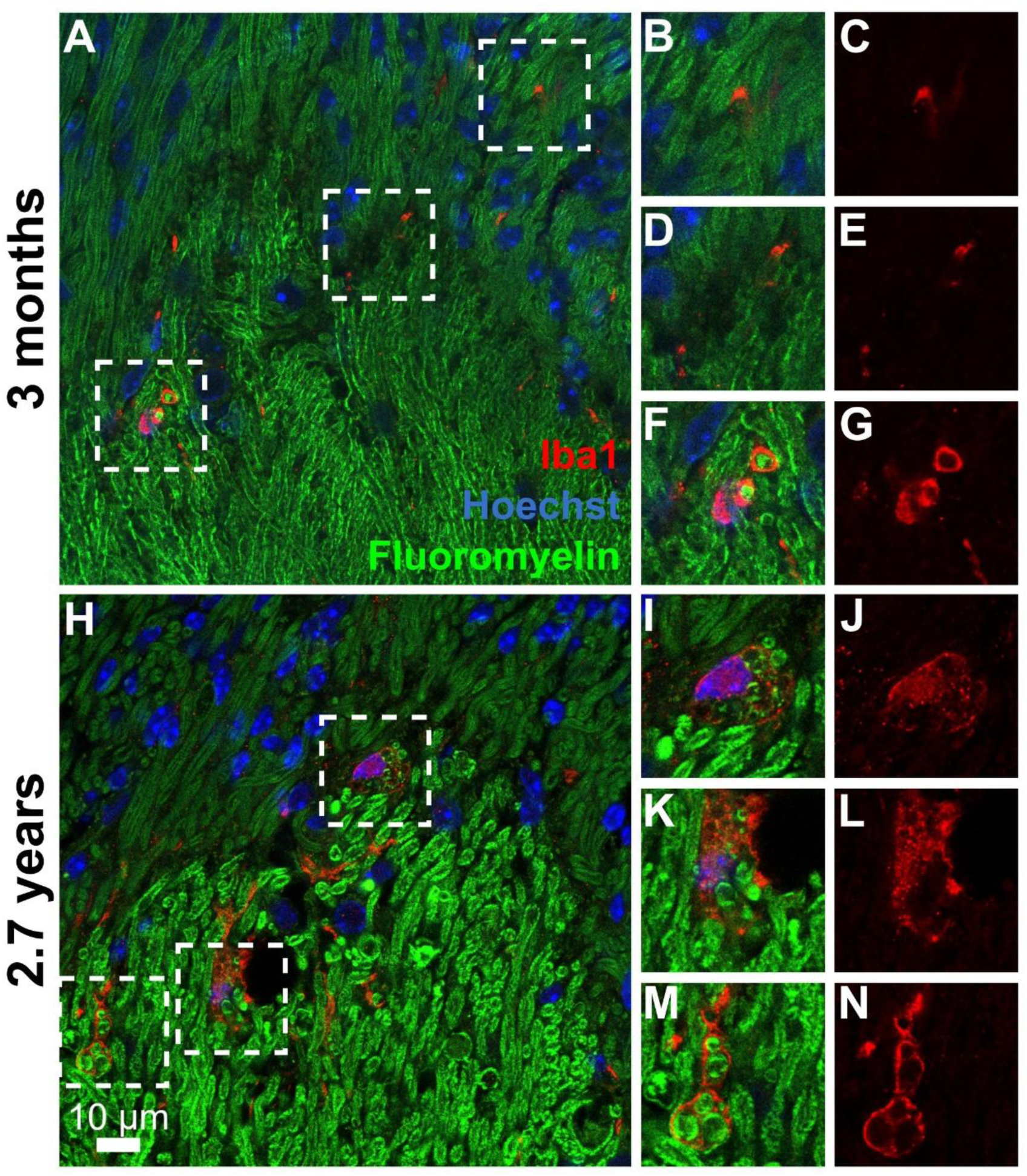
Increased healthy-like myelin found in Iba1^+^ cells in the aging AN GTZ. **(A-N)** Sections of the AN GTZ in young (A-G) and old (H-N) mice showing Iba1^+^ cells with internalized myelin. Dotted boxes in A and H are regions that were enlarged to show the structurally intact myelin within Iba1^+^ macrophages/microglia at the AN GTZ (B-G, I-N). Images are individual slices from the z-stack to show the structurally intact myelin within Iba1^+^ cells.

### 3.7 Human temporal bones have Iba1^+^ cells with internalized myelin at the auditory nerve glial transition zone

To further validate the age-related changes in the immune-glial interactions determined in mice, we analyzed human temporal bone sections at the AN GTZ. We used sections of the temporal bones from an aged donor (89+ years old) to look at aging effects on this region of the AN. Locating mid-modiolar sections from human temporal bones is challenging, and identifying the specific sections that contain the GTZ is difficult due to the large number of sections (>200 per specimen) and the exact angle required to capture the AN GTZ. Once we located the appropriate human temporal bone sections, sections were immunostained and then imaged to evaluate the Iba1^+^ macrophages/microglia present at the AN GTZ (Figure 8). We identified Iba1^+^ cells throughout the human AN. The GTZ was again identified in human sections based on the difference in density of nuclear staining (more nuclei peripherally due to myelination by Schwann cells and fewer nuclei centrally due to myelination by oligodendrocytes) (Figure 8A). Additionally, we often found Iba1^+^ cells around the AN GTZ, similar to the mouse AN GTZ. High-resolution imaging revealed Iba1^+^ cells in direct contact with myelin sheaths of the AN both centrally and peripherally (Figure 8B). Iba1^+^ cells containing internalized myelin debris were also observed (Figure 8B, C), perhaps suggesting phagocytic processes occur in the aging human AN and are amplified at the AN GTZ.

**Figure 8.**
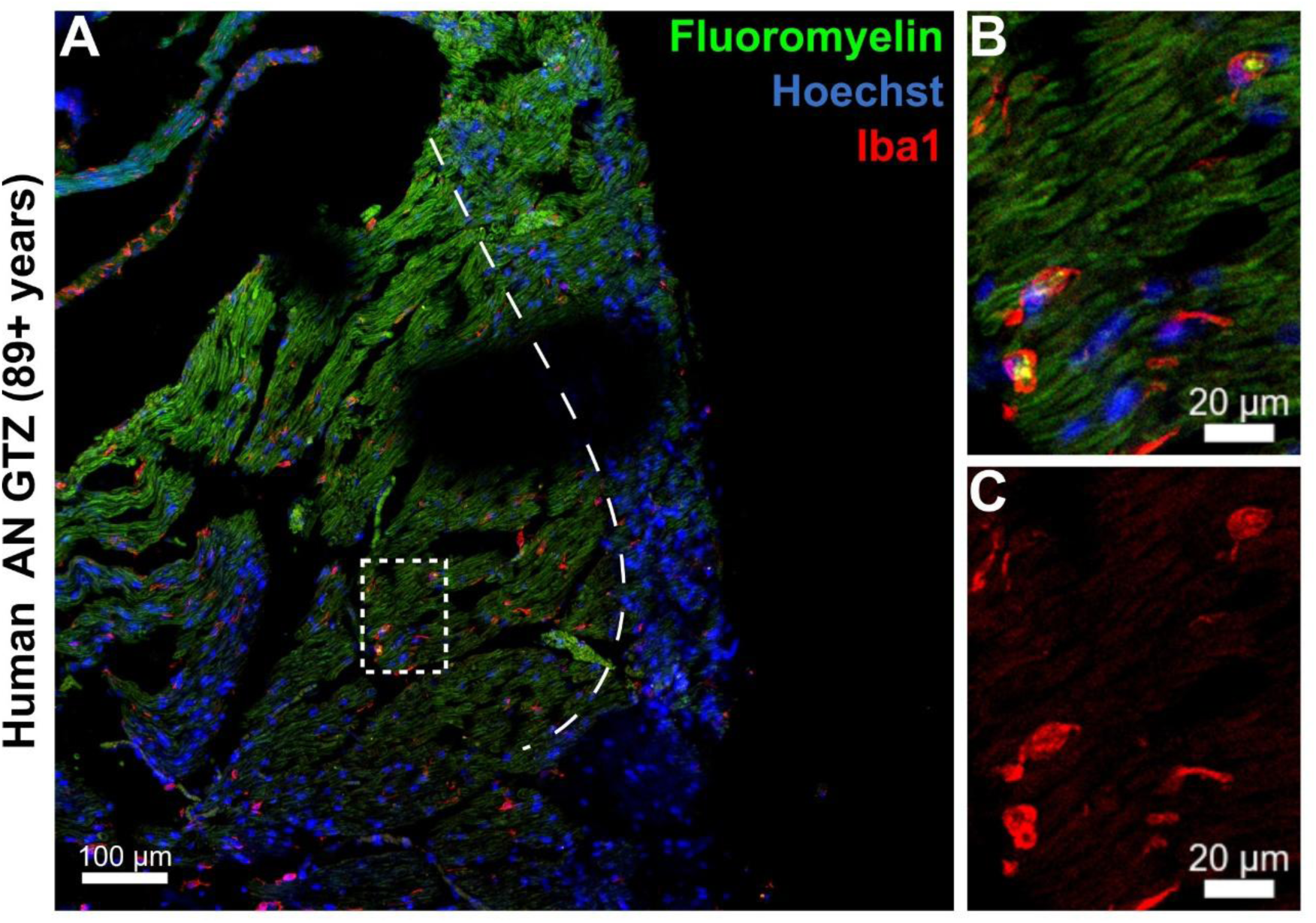
Internalized myelin found in human AN Iba1^+^ cells. **(A)** Mid-modiolar cochlear section containing the AN GTZ in a temporal bone from an 89+ year old donor. Large white dashed line is where the AN GTZ is located. **(B, C)** Enlarged images from A showing Iba1^+^ cells in close association with AN myelin and containing internalized myelin throughout the AN. Dashed box in A is shown in B and C. Sections are stained with FluoroMyelin^TM^ (green), Iba1 (red), and Hoechst (blue). Scale bar = 100 µm.

## 4. Discussion

The present study identifies the AN GTZ as a dynamic neuroimmune interface with age-related structural changes that may play a role in AN myelin degeneration and functional declines associated with ARHL. By integrating quantitative immunohistochemistry, morphological analyses, 3D high-resolution imaging, and transcriptomic profiling, we demonstrate that this central-peripheral interface at the AN GTZ is not just a boundary between Schwann cells and oligodendrocytes, but a dynamic region characterized by enhanced immune activation and immune-glial interactions in the aging AN. Aging was associated with reduced myelination and disrupted peripheral and central myelin sheaths as well as a substantial increase in Iba1^+^ macrophages/microglia throughout the AN, with significant enrichment of abnormal immune activity at the GTZ. This heightened immune presence at the AN GTZ suggests enhanced focal vulnerability to immune dysregulation contributing to impaired myelin homeostasis and AN degeneration. Validation of mouse AN observations in human temporal bones from older donors further supports the abnormal phagocytic activities by immune cells within the AN and its GTZ, highlighting demyelination and enhanced glial dysfunction as key contributors to AN degeneration and ARHL. Together, these findings position the GTZ as a unique nexus of neuroimmune activity with broad implications, not just for understanding and treating AN degeneration, but also for better understanding the specialized biology of GTZs throughout the central and peripheral nervous systems.

### 4.1 A unique central-peripheral immune interface at the auditory nerve glial transition zone

The AN connects the peripheral sensory epithelia housed within the cochlea to the central auditory pathways, yet the glial architecture and immune microenvironment at its central-peripheral transition remains largely unexplored. Previous studies noted the existence of GTZ regions and their development across spinal and cranial nerves (Moll and Meier, 1983; Fraher, 1992; Fraher and Cheong, 1995; Toma, McPhail and Ramer, 2006; Ziyal and Ozgen, 2007). In the auditory system, coordinated migration of oligodendrocytes at the AN GTZ during development creates a border between peripheral and central in cat, rat, and mouse models (Knipper *et al*., 1998; Osen, Furness and Hackney, 2011; Bojrab *et al*., 2017).

Because this region represents a structural and immunological boundary, we examined how immune cells and myelinating glia interact at this unique central-peripheral interface at the AN GTZ. It is well established that macrophages are required for normal auditory function, but can also contribute to the pathogenesis of injury and disease (Sung *et al*., 2019, 2024; Chokr *et al*., 2022; Seicol, Lin and Xie, 2022; Shimada *et al*., 2023). Traditionally, the AN studies focused on either the peripheral AN, maintained by macrophages and Schwann cells, or the central auditory system and cochlear nucleus, maintained by microglia and oligodendrocytes (Wan and Corfas, 2017; Long *et al*., 2018; Xin *et al*., 2025). Numerous studies report cochlear macrophage activity during development and their roles in spiral ganglion neuron health after hair cell injury and noise exposure (Hirose *et al*., 2005; Shi, 2010; Kaur *et al*., 2015; Hirose, Rutherford and Warchol, 2017; Dong *et al*., 2018; Kishimoto *et al*., 2019; Miwa *et al*., 2024; Murali *et al*., 2025). The role of microglia has been investigated in the central auditory system, including the cochlear nucleus and auditory brainstem during development and after injury (Baizer *et al*., 2015; Cramer and Rubel, 2016; Fuentes-Santamaría *et al*., 2017; Milinkeviciute and Cramer, 2018; Milinkeviciute *et al*., 2019; Wang *et al*., 2020). Here, our study offers a new approach for studying the cochlear macrophage/microglia populations and their immune-glial interactions across both the central and peripheral nervous systems, both of which are contained within the cochlear modiolus.

Our data revealed that the AN GTZ contains a sharply demarcated shift in myelination between the peripheral and central AN regions accompanied by a myeloid compartment (macrophages/microglia) that is far less segregated in the GTZ than assumed, based on the canonical understanding of the central and peripheral nervous systems (Figure 1). At the AN GTZ the processes of some Iba1^+^ cells extend to both the peripheral and central AN (Figure 3), suggesting that macrophages/microglia move freely across this boundary, and exhibit modification in their phagocytic activity specifically within the GTZ, indicated by the numerous Iba1^+^ cells with internalized healthy-like myelin found only at the GTZ with aging (Figure 7).

This central-peripheral interface located within the cochlear modiolus also represents a potential immunological niche where macrophages and microglia coexist and may interact to coordinate their responses to pathologic conditions, such as aging. Notably, there is no physical barrier separating the peripheral and central immune microenvironments; rather, the GTZ is defined by a discrete gap in myelination where the shift between Schwann cells and oligodendrocytes occurs (Fraher, 1992; Bojrab *et al*., 2017). One study reported that glia can adapt and change their phenotype in response to dysfunctional neighboring glia (Beachum *et al*., 2025). It has also been reported that microglia mount a specialized response at root entry zones in both the peripheral and central portions of the sciatic nerve after injury (Gai, Zhou and Rush, 1996; Liu, Rudin and Kozlova, 2000). Additionally, studies have reported that the various glial cells, including oligodendrocytes, astrocytes, Schwann cells, and macrophages/microglia, have plasticity in phenotype in response to injury (Luo *et al*., 2019; Chokr, Bui-Tran and Cramer, 2024). In our study, the AN GTZ displays region-specific pathological changes with aging, including a marked increase in macrophage/microglia density, suggesting that the GTZ may serve as a specialized site of immune surveillance and signaling distinct from the surveillance processes that occur in the central or peripheral compartments.

### 4.2 Abnormal phagocytic activity may be a key contributor to age-associated demyelination of the auditory nerve

A defining feature of the aging AN in our study was the coordinated emergence of demyelination and impaired phagocytic activity in the AN, with evidence of immune system dysregulation at the AN GTZ. It is well established that aging is associated with demyelination in the brain (Réu *et al*., 2017; Hill, Li and Grutzendler, 2018; Hou *et al*., 2023). Additionally, aging has been linked with phagocytic dysfunction in the brain, especially in the context of neurodegenerative disease, some of which can cause hearing loss (Leszek *et al*., 2016; Raj *et al*., 2017; Zheng *et al*., 2017; Colonna and Brioschi, 2020; Butler *et al*., 2021; Beirowski, 2022, 2022; Silvin *et al*., 2022; Huang *et al*., 2025). Our data showed that aging is associated with a reduction in myelin integrity throughout the peripheral and central AN, reflected by a disruption in myelin sheaths and an accumulation of myelin debris within macrophages/microglia (Figures 4, 6, and 7). This is particularly interesting because previous studies in the brain have reported that myelin-loaded microglia is evidence of phagocytic dysfunction, characteristic of a proinflammatory phenotype, and can exacerbate disease progression (O’Neil *et al*., 2018; Thomas *et al*., 2022; Quick *et al*., 2023; Gao *et al*., 2024). However, future studies should focus on identifying specific macrophage and microglia phenotypes and their phagocytic function that may be contributing to AN degeneration and should use advanced omics and direct assay of phagocytic activity in the AN. This convergence of reduced myelination throughout the AN and evidence of proinflammatory phagocytic dysfunction in the Iba1^+^ cell population suggests that the GTZ may be a focal point for age-related myelin degeneration.

Although the presence of Iba1^+^ cells in the cochlea and AN has been established, their function in AN myelination and aging remains unclear (Defourny, Lallemend and Malgrange, 2011; Ginhoux and Guilliams, 2016; Bojrab *et al*., 2017; Brown *et al*., 2017; Driver and Kelley, 2020; Yu, Gao and Wan, 2021; Hough *et al*., 2022). Myelination is an ongoing process that occurs throughout a person’s lifetime (Bacmeister *et al*., 2020; Langley, Triplet and Scarisbrick, 2020; Franklin, Frisén and Lyons, 2021; Sen *et al*., 2022; Kent and Miron, 2023). The roles of macrophages/microglia in myelin maintenance have been reported in both the peripheral and central nervous systems (Koike and Katsuno, 2021; Kopper *et al*., 2021; Ryan *et al*., 2022; Sen *et al*., 2022). In the aging brain, deficits in phagocytic activity associated with myelin turnover and maintenance are often found in neurodegenerative and demyelinating diseases such as Alzheimer’s Disease, Multiple Sclerosis, and Amyotrophic Lateral Sclerosis (Bitsch *et al*., 2000; Cunha *et al*., 2020; Gaitsch *et al*., 2024). Our morphological analyses revealed that Iba1^+^ macrophages/microglia in aged mice often contained myelin debris, suggesting active but incomplete phagocytic engagement (Figures 6 and 7). Together with transcriptomic analysis of aged ANs, our data suggests a decline in effective phagocytosis, with aged AN Iba1^+^ cells exhibiting signs of stalled or dysfunctional clearance rather than efficient debris removal. This dysfunctional clearance may be linked to the upregulation of *Clec7a*, *Trem2*, and *Itgb2* (Figure 6). These genes have been linked to dysregulation of phagocytic processes and are expressed by disease-associated microglia in the brain (Jay, Von Saucken and Landreth, 2017; Deczkowska *et al*., 2018; Deerhake and Shinohara, 2021; Silvin *et al*., 2022; Gao *et al*., 2023; Martins-Ferreira *et al*., 2025). However, our study utilized bulk RNA sequencing of the ANs which does not allow regional resolution between the central AN, peripheral AN, and GTZ. Future studies should focus on heterogeneity of the Iba1^+^ cell populations in the AN and cochlea using advanced omics approaches such as single-cell RNA sequencing and spatial omics assays to better define the spatial organization of age-related changes of AN immune cells.

In addition, lipid-burdened macrophages/microglia can take on a proinflammatory state and exacerbate disease progression (Burns *et al*., 2020; Franklin and Simons, 2022). Additionally, when myelination is disrupted, this can lead to axonal damage and neuronal cell death (Beirowski, 2022). This impaired phagocytic function likely contributes to the persistence of myelin debris observed throughout the aged AN and may exacerbate degeneration by failing to restore a supportive microenvironment for AN axon health and conduction (Sepp, Schulte and Auld, 2001; Beirowski *et al*., 2014, 2014; Leszek *et al*., 2016; Arancibia-Cárcamo *et al*., 2017; Beirowski, 2022, 2022; Grüter *et al*., 2022; Freire *et al*., 2023).

More importantly, this decrease in myelination was not restricted to either canonical peripheral or central AN compartment. Both the Schwann cell-associated and oligodendrocyte-associated regions exhibited signs of compromised myelin turnover or repair (Figure 4), indicating that aging induces a similar maladaptive immune-glial response across the entire AN, with enhanced effects at the GTZ. Together, our findings support a model in which aging leads to widespread demyelination coupled with a dysfunctional immune cell compartment with insufficient AN self-repair mechanisms, culminating in progressive AN degeneration and ARHL (Figure 9).

**Figure 9.**
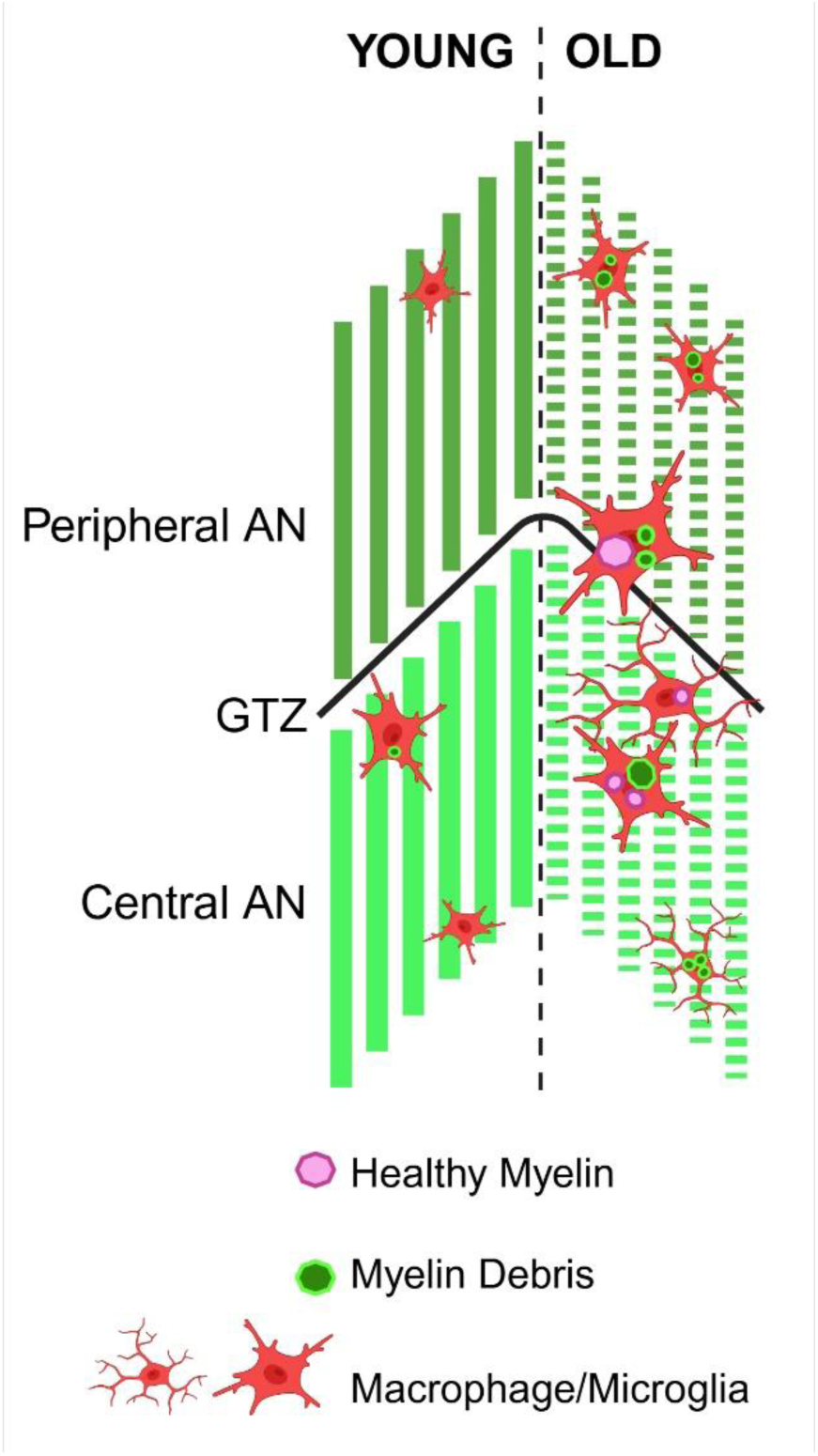
Summary of Aging Effects on the AN and GTZ. Schematic demonstrating age-related decreased myelination, increased numbers of Iba1^+^ macrophages/microglia, and enhanced dysfunction at the AN GTZ. The model highlights regional differences in age-related immune and glial changes across the peripheral AN, central AN, and GTZ compared to the young AN. In the aging AN, Iba1^+^ cells throughout all regions of the AN contain myelin debris, while the GTZ exhibits a marked increase in Iba1^+^ cells that also contain structurally intact myelin, suggesting region-specific functional declines that may contribute to age-related AN demyelination.

### 4.3 A dynamic glial transition zone is conserved in the human auditory nerve

To extend our studies from the mouse AN to humans, we examined the structure and immune composition of the GTZ in the human AN from an 89+ year old donor, revealing a similarly dynamic region with enriched immune-glial interactions. The GTZ of the AN is a unique neuroanatomical boundary where central and peripheral myelination converge, which we observe in our study in human temporal bones (Skinner, 1931; Guclu *et al*., 2009). Our study confirmed that an active myeloid population is present in the human AN GTZ. Another important observation was that Iba1^+^ cells were identified containing internalized myelin debris (Figure 8), suggesting that phagocytic processes also occur in humans to support myelin homeostasis and normal function of the AN. It has been shown that microglia turnover in the human brain is a slow continuous process, making this slowly renewing population particularly vulnerable to pathologic disturbances like aging and demyelination (Réu *et al*., 2017). Additionally, the unique biology of the AN GTZ region may be a contributor to the increased rates of schwannoma in the CNVIII compared to other cranial nerves, as well as a driver of dysfunction in these glia leading to the development of schwannomas across the nervous system (Nickele *et al*., 2012; Lan *et al*., 2020; Eggink *et al*., 2022). Some studies have linked variation in central myelin morphology in the trigeminal nerve to the development of trigeminal neuralgia (Nomura *et al*., 2019). Additionally, studies are focusing on this region due to its regenerative potential in the context of spinal cord injury (Carlstedt, 1997; Monje, Deng and Xu, 2021; Ghosh and Pearse, 2023). The localization of Iba1^+^ cells to this region and the evidence of phagocytic activity point to potential roles in myelin maintenance and remodeling. These processes are critical to AN function and any dysregulation could contribute to age-related demyelination, AN degeneration, and ARHL.

Our findings raise the possibility that targeting immune-glial interactions at the GTZ could be a novel strategy to mitigate age-related demyelination in the AN. Preclinical models have shown that modulating glial subtype can have a protective role, particularly in neurodegenerative disease (Rahimian *et al*., 2022; Silvin *et al*., 2022; Barclay *et al*., 2023; Gao *et al*., 2023; Ball *et al*., 2024; Martins-Ferreira *et al*., 2025). Modulating macrophage/microglia activation state to restore phagocytic function may help maintain myelin integrity and preserve axonal conduction in aging. Interventions aimed at supporting this dynamic immune niche at the GTZ could be potential strategies to ultimately slow or prevent the progression of AN degeneration and ARHL.

## Data Availability Statement

The original contributions presented in this study are included in the article; further inquiries can be sent to the corresponding authors.

## Ethics Statement

All animal studies for this project were reviewed and approved by the Institutional Animal Care and Use Committee of the Medical University of South Carolina (MUSC) and the Ralph H. Johnson VA Health Care System (VAHCS) in Charleston, South Carolina.

## Author Contributions

SP and HL designed the study. HL, SP, PC, and HY acquired the funding. SP, HA, and HL prepared samples. SP performed confocal imaging. PC, JC, and HY developed and performed light sheet imaging. JB analyzed the RNA-seq dataset. SP developed unpublished analysis routines. SP analyzed the data. SP wrote the manuscript. SP, HL, and HA revised the manuscript. All authors contributed to the article and approved the submitted version.

## Acknowledgements

We thank Jiaying Wu for her technical assistance and Tyreek Jenkins and Emily Fabrizio-Stover for their comments and suggestions.

