## Supplemental Figures for "A Novel Central-Peripheral Interface: The Auditory Nerve Glial Transition Zone Exhibits Enhanced Age-Related Immune and Glial Cell Dysfunction"

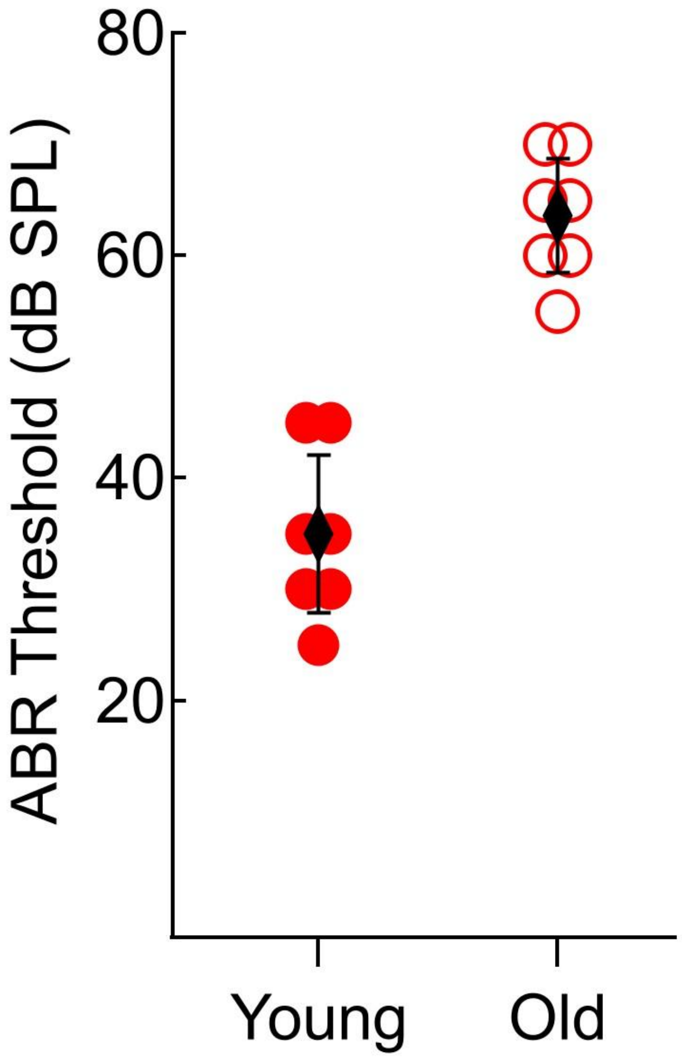


**Supplementary Figure 1. Auditory brainstem response (ABR) thresholds of CBA/CaJ mice used in this study.** ABR thresholds at 11.3 kHz for young (3-4 months) and old (2.5+ years) mice of both sexes used in this study. Red markers indicated the young individuals (red filled circles; n=7) and old individuals (red open circles; n=7). Means are indicated by black diamonds. Error bars indicate the standard deviations for each group.

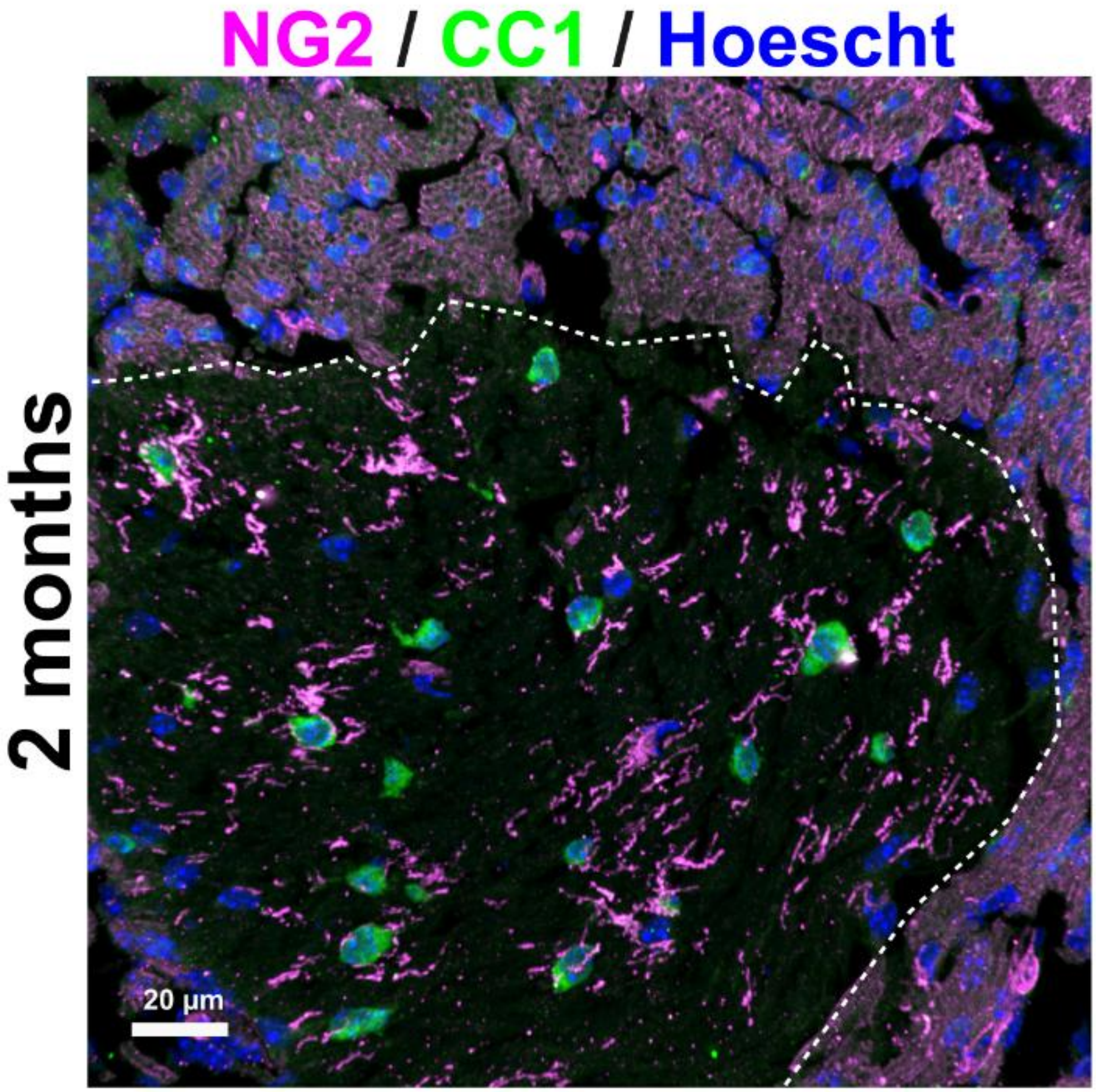


**Supplementary Figure 2. Oligodendrocyte markers to differentiate the central AN from the peripheral AN.** Representative image of the AN GTZ in a young (2 months) CBA/CaJ mouse. CC1, a marker for mature oligodendrocytes that does not label myelin, is shown in green. NG2, a marker for oligodendrocyte progenitor cells (OPCs), is shown in magenta. In the AN, NG2 stains OPCs in the central AN, but also Schwann cells populating the peripheral AN. The white dashed line indicates the GTZ.
